# Improving the accuracy of cerebral blood flow measurement by phase contrast MRI

**DOI:** 10.1101/2024.08.13.607816

**Authors:** Xiuli Yang

## Abstract

Cerebral blood flow (CBF) is a critical hemodynamic marker for natural aging and pathological conditions. It can be assessed non-invasively by the phase-contrast (PC) magnetic resonance imaging (MRI) technique. Although the technical principle of PC MRI is straightforward, related experimental settings, e.g., spatial resolution, partial volume effect, slice positioning angle, and signal-to-noise ratio (SNR), require deliberate considerations to ensure measurement accuracy. In this study, we performed simulations to systematically investigate the dependence of measurement accuracy of PC MRI on the spatial resolution, extent of partial volume effect, slice imperfection angle, and SNR. It has been found that at least 6 voxels along the vessel diameter are required to ensure accurate CBF measurements. Partial volume effect acts to underestimate CBF measurements. The tolerance against slice imperfection angle is ≤15 degree for the commonly seen artery in mice under isoflurane anesthesia. A normalized SNR of 25.47 dB is required to ensure the accuracy of CBF measurement. Our study will promote the utilization of CBF as a pathophysiological marker for future studies by delineating the factors affecting measurement accuracy in PC MRI.

## Introduction

The past few decades have witnessed the ubiquitous methodological development and applications of magnetic resonance (MR) technique in various fields ^1-5^. MR technique is associated with excellent versatility typically achieved via elaborate designs of radiofrequency pulses, pulsed field gradients, and acquisitions, thus providing a variety of spatial or spectral information ^6-9^. Based on this versatility, MR technique allows multiparametric mapping in a same MRI session, avoiding the necessity of co-registration across different modalities and prolonged experimental preparations of several different techniques.

Functional magnetic resonance imaging (fMRI) has attracted broad research interests since its advent by providing real-time functional responses to designed stimulations, unraveling the neuronal functional connectivity, and estimating static or dynamic physiological parameters ^10-14^. Bold-oxygenation-level-dependent (BOLD) MRI is the most popularly used fMRI method ^15,16^. However, it suffers from the disadvantage of multifaceted signal contributions from local blood oxygenation, hematocrit level, cerebral blood volume, and cerebral blood flow (CBF) ^17^. Therefore, paralleling the technical development within the framework of BOLD, MRI sequences focusing on a specific physiological parameter have been actively developed, for example, T2-relaxation-under-spin-taggging (TRUST) for measuring venous oxygenation ^18-20^, phase contrast (PC) for measuring total CBF ^21,22^, arterial-spin-labeling (ASL) for measuring regional perfusion ^23^, vasoactive challenge based dynamic MRI for measuring cerebrovascular reactivity (CVR) ^24-26^, and ASL based modeling for evaluating blood-brain-barrier (BBB) function ^27-29^. These novel techniques not only provide quantitative evaluations on a specific function, but also facilitate the interpretation of BOLD signals.

CBF plays a central role in the microvascular function. As revealed by BOLD, CBF increase is a primary response to a neuronal stimulation for supplying the locally increased metabolic demand. PC MRI utilizes a pair of bipolar gradients (with equal first momenta) to encode flow information by sensitizing its phases to blood velocities ^22^. With the velocity map, blood flow within a selected artery can be obtained by the integral of velocities over the arterial voxels. By covering the major feeding arteries of brain ^30^, i.e., left/right internal carotid arteries (LICA/RICA) and left/right vertebral arteries (LVA/RVA), total blood flow can be calculated and further converted into CBF in the unit of ml/100g/min after accounting for the brain weight. Since LVA and RVA merge into the basilar artery (BA), BA is often observed as an alternative when targeting at the total blood flow. The performance of PC MRI heavily depends on the experimental setup, e.g., spatial resolution, partial volume effect, slice positioning angle, and signal-to-noise ratio (SNR). In this study, we employed simulation to investigate the dependence of measurement accuracy on these experimental factors.

## Methods

### Simulation

The flow pattern in arteries was assumed to be laminar, i.e., 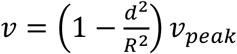, where *d* denoted the distance by reference to the vessel center, *R* the vessel radius, and *v*_*peak*_ the peak velocity. Using this equation, blood flow at the vessel center was assumed to be peak velocity and flow at the vessel wall was assumed to be 0 cm/s. Mouse was used as an example to make assumptions. Velocity map was calculated for a field of view (FOV) of 2000×2000 µm2 and a simulation resolution of 1 µm. Velocity map was used to calculate phase under the encoding velocity (VENC) of 20 cm/s. All simulation scripts were written based on the MATLAB platform.

### Study 1: Dependence of CBF accuracy on the spatial resolution

Peak velocity was assumed at 15 cm/s and partial volume effect was 1/4 (i.e., the intensity ratio between a voxel with pure tissue and a voxel with pure blood). A total number of 7 different vessel diameters were simulated: 100, 200, 300, 350, 400, 450, and 500 µm. Acquisitions were simulated under a series of spatial resolutions from 1 µm to 100 µm at the step-wise interval of 1 µm. Slice positioning of PC MRI was assumed to be perfectly perpendicular to the vessel trajectory.

### Study 2: Dependence of CBF accuracy on the extent of partial volume effect

Peak velocity was assumed at 15 cm/s and vessel diameter was 300 µm. A total number of 7 different extents of partial volume effect was examined: 0, 1/10, 1/8, 1/6, 1/4, 1/2, and 1. The influence of partial volume effect was simulated under a series of spatial resolution (1-100 µm with the step-wise interval of 1 µm). Slice positioning of PC MRI was assumed to be perfectly perpendicular to the vessel trajectory.

### Study 3: Dependence of CBF accuracy on the imperfection of slice positioning

Peak velocity was assumed at 15 cm/s, spatial resolution was 50 µm, and partial volume effect was assumed to be 1/2. Six vessel diameters were simulated, including 200, 300, 350, 400, 450, and 500 µm. Slice thickness was assumed to be 500 µm and the simulation resolution for slice thickness was 50 µm. The slice positioning imperfection was represented by the titled angle by reference to the perpendicular case (i.e., perpendicular case will be denoted as slice imperfection angle of 0°). The imperfection angles from 0° to 35° were simulated at the step-wise interval of 1°.

### Study 4: Dependence of CBF accuracy on SNR

Peak velocity was assumed at 15 cm/s and partial volume effect was assumed to be 1/2. Six vessel diameters were simulated, i.e., 200, 300, 350, 400, 450, and 500 µm. Slice thickness was assumed to be 500 µm and a basic simulation element had a thickness of 50 µm. Slice imperfection angle was assumed to be 0°. A series of normalized noise levels from 0% to 15% was simulated at the step-wise interval of 1% (assuming blood flow was 100%). Each data point was simulated for 10 repetitions.

### Statistical analyses

Linear regression analyses were employed to examine dependence of BF accuracy on different experimental factors. Spearman’s correlation was utilized to examine the associations between optimal settings and the investigated factors. P<0.05 was considered significant.

## Results

### Study 1: Dependence of CBF accuracy on the spatial resolution

According to the linear regression, BF exhibited positive dependences on the spatial resolution (P<0.001, Figure 1A) and vessel diameter (P<0.001, Figure 1A). The diameter dependence simply meant that larger arteries had larger BF (equal peak velocity assumed). The resolution dependence indicated that accurate BF measurement required sufficiently higher spatial resolution. Otherwise, there was a BF overestimation. Similar results can be found in the normalized BF (Figure 1B). The measured peak velocity exhibited a negative dependence on the spatial resolution (P<0.001, Figure 1C), indicating that high spatial resolution is desired for the accurate estimation of peak velocity. Meanwhile, peak velocity was associated with a positive diameter effect primarily driven by the resolution effect. In other words, when spatial resolutions were sufficiently high, the measured peak velocities were not different across vessel diameters. If 5% error was allowed for the normalized BF (Figure 1B), we could obtain the optimal spatial resolution for different vessel diameters. There was a good correlation between optimal resolution and vessel diameter (R2=0.999, P<0.001, Figure 1D). Based on the correlation equation, it could be calculated that the least voxel number along the vessel diameter to avoid overestimation should be 6, i.e., 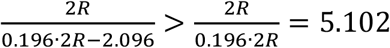.

**Figure 1.**
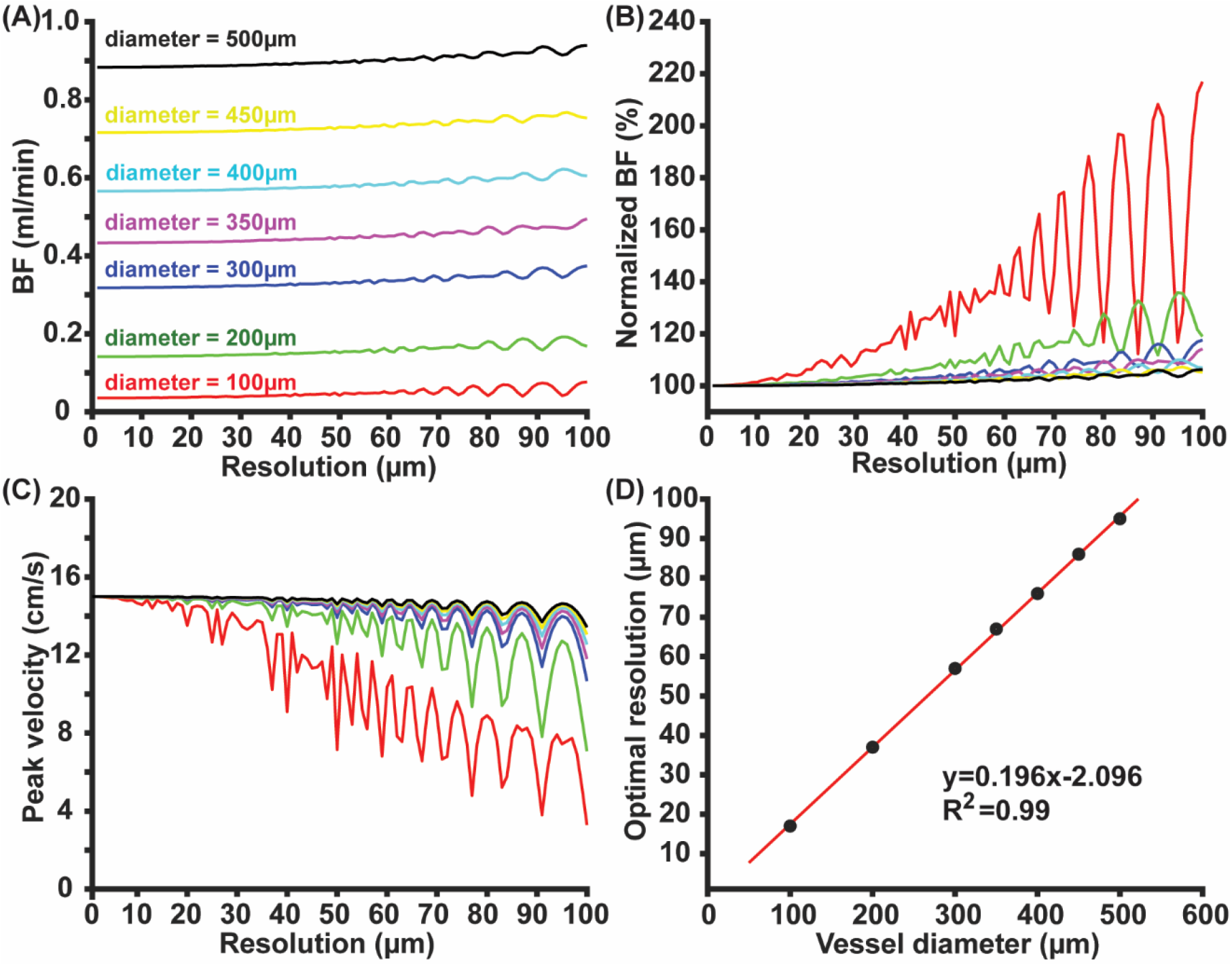
Dependence of BF measurement on spatial resolution. (A) BF as functions of spatial resolutions; (B) normalized BF as functions of spatial resolutions; (C) peak velocity as functions of spatial resolution; (D) correlation between the optimal resolution and vessel diameter.

### Study 2: Dependence of CBF accuracy on the extent of partial volume effect

The BF was associated with a positive dependence on spatial resolution (P<0.001, Figure 2A) and a negative dependence on partial volume effect (P<0.001, Figure 2A). These two factors counteracted each other. As a result, the overestimation induced by insufficient spatial resolutions became less prominent in the presence of a relatively stronger partial volume effect (particularly, for PVE=1 in Figure 2A). Similar results could be found in the normalized BF (Figure 2B). In contrast, peak velocity was not affected by the partial volume effect (Figure 2C) due to the fact that partial volume effect merely happened at the boundary region. If 5% error was allowed in the normalized BF, there was a significant positive correlation between optimal spatial resolution and normalized partial volume effect (R2=0.974, P<0.001, Figure 2D). Namely, when there was a stronger partial volume effect, the requirement on spatial resolution became looser.

**Figure 2.**
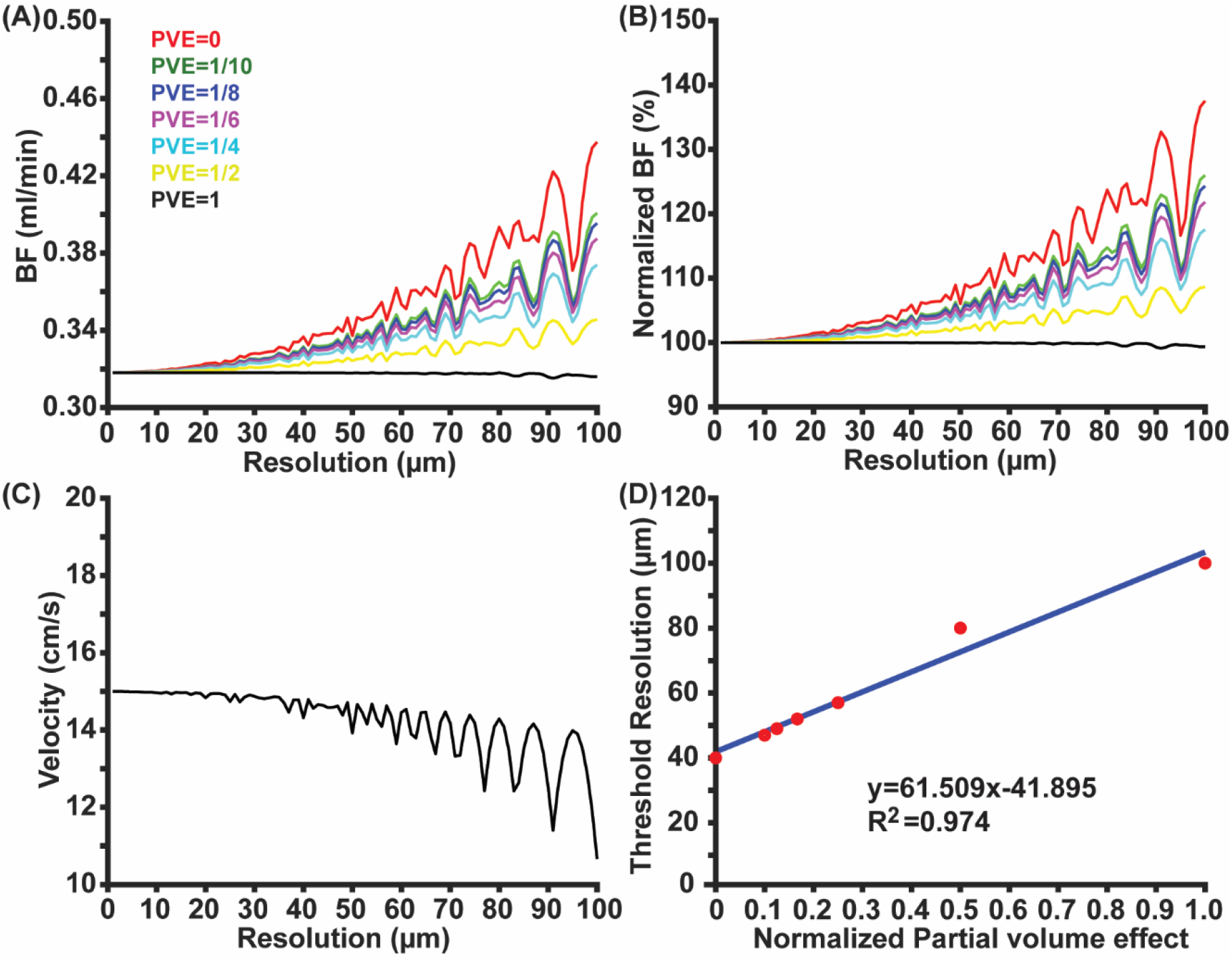
Dependence of BF measurement on the partial volume effect. BF (A), normalized BF (B), and peak velocity (C) as functions of spatial resolution under different partial volume effect. In (C), only the curve of PVE=1 could be visualized because all curves overlapped with each other. (D) correlation between the optimal resolution and normalized partial volume effect.

### Study 3: Dependence of CBF accuracy on the imperfection of slice positioning

BF exhibited a positive dependence on the slice imperfection angle (P<0.001, Figure 3A), indicating that improper slice positioning led to overestimation. Such a trend was more obvious in the normalized blood flow (Figure 3B). There was a positive correlation between the tolerable slice imperfection angle and vessel diameter (R2=0.777, P=0.02, Figure 3C), indicating that BF measurement of larger arteries held stronger tolerance against the slice imperfection angle. One possible explanation was that larger arteries had broader circumstances under imperfect slice positioning and thereafter more partial volume effect to counteract the overestimation.

**Figure 3.**
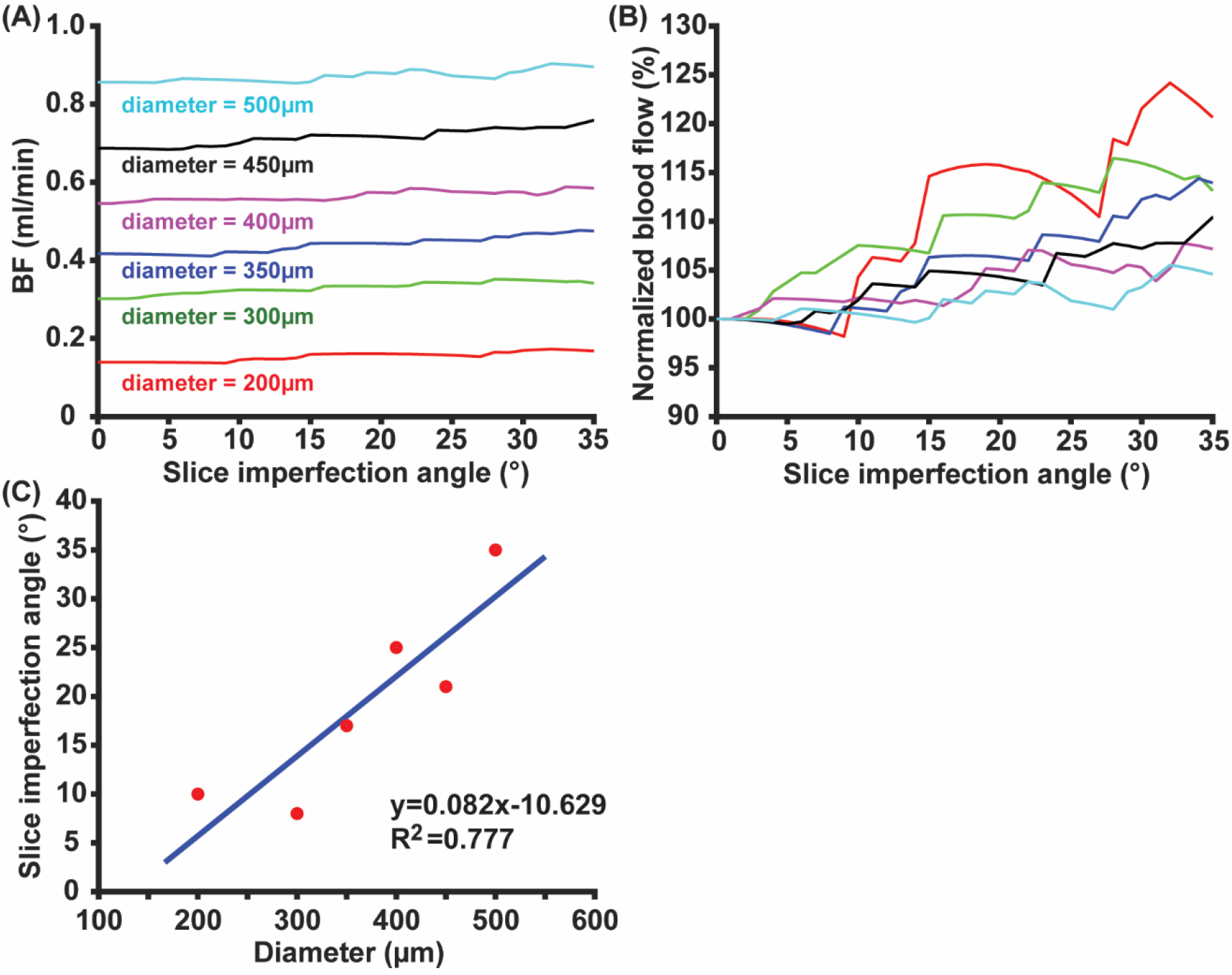
Dependence of BF measurement on slice positioning. (A) and (B) denote BF and normalized BF as functions of slice imperfection angle; (C) correlation between the tolerable slice imperfection angle and the vessel diameter.

### Study 4: Dependence of CBF accuracy on SNR

Across the different vessel diameters, BF tended to be less stable at larger noises (Figure 4A-4F), which was well within theoretical expectations. If a coefficient-of-variation (CoV) of 5% was allowed, the tolerable normalized noise level was 5.33±0.82% (Mean ± Standard deviation), corresponding to 25.47 decibels.

**Figure 4.**
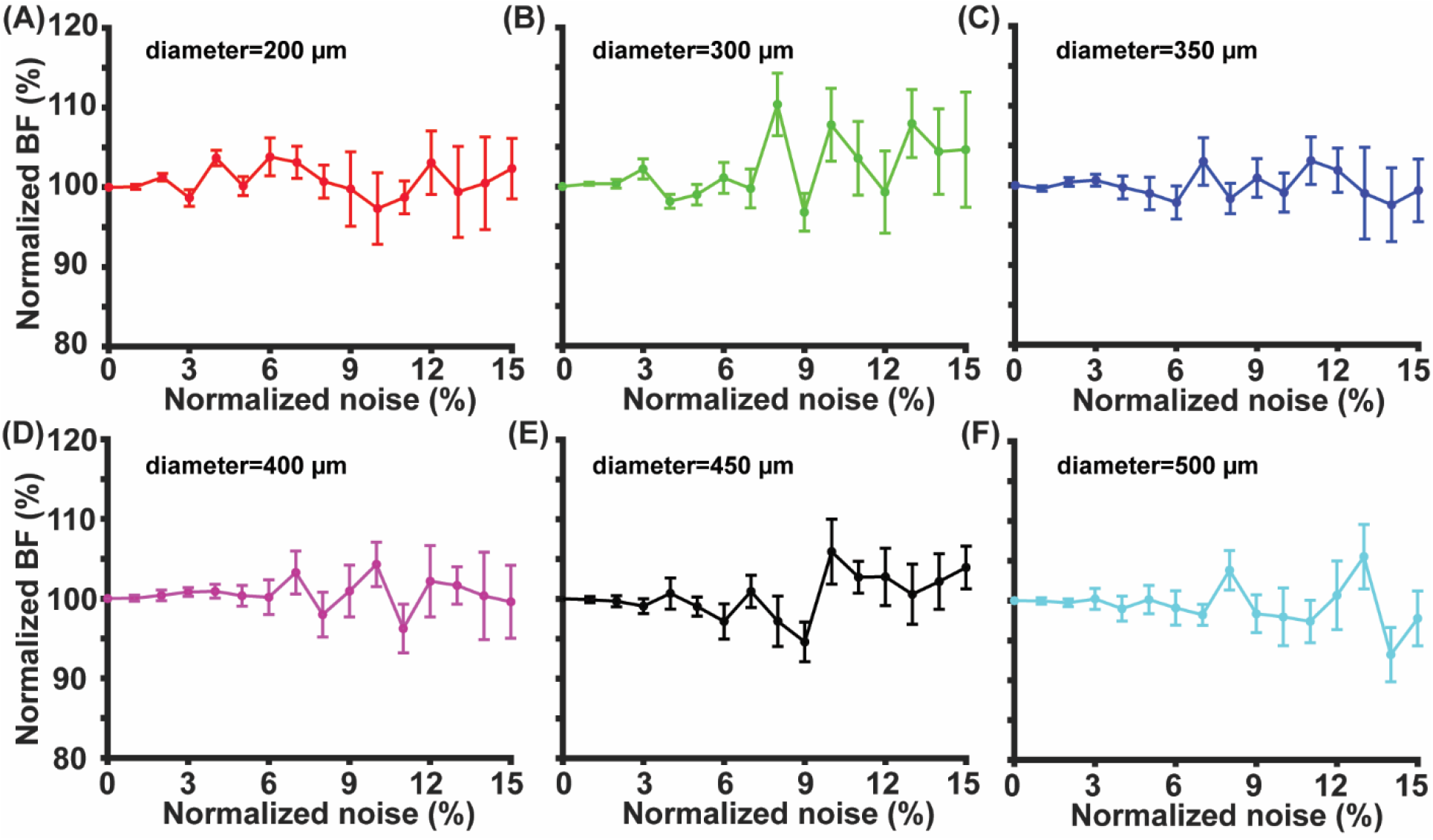
Dependence of BF measurement on the normalized noise level under the diameter of 200 (A), 300 (B), 350 (C), 400 (D), 450 (E), and 500 (F) µm. Error bar represented the standard error over 10 simulation repetitions.

## Discussion

We have systematically investigated the influence of critical parameters in PC MRI on the measurement accuracy. The optimal spatial resolution depends on the arterial diameter and at least six voxels along the vessel diameter is the boundary resolution to ensure accuracy (5% error allowed). Contrary to the overestimation induced by insufficient spatial resolution, partial volume effect will underestimate the CBF measurement. Slice orientation imperfection can induce CBF overestimation and the maximal tolerable imperfection angle is positively correlated with the vessel diameter. A normalized noise level no larger than 5.3% is required for accurate CBF measurement.

In an optimization study of mouse PC MRI, the diameter of LICA is approximately 338 µm in mice and optimal spatial resolution is 50 µm ^21^. Our conclusion that at least 6 voxels along the vessel diameter is required to ensure measurement accuracy shows agreement with this experimental data. Spatial resolution can be sharpened by increasing the matrix size while preserving FOV at the cost of prolonged acquisition time.

The strength of partial volume effect can be modulated by repetition time (TR). When a shorter TR is used to improve the acquisition efficiency, tissue signal will be smaller due to less longitudinal relaxation recovery. SNR can be enhanced by utilizing state-of-the-art probes (e.g., cryoprobe) or adding signal averages ^31^. In the application of PC MRI, a short TR is often used due to the short apparent longitudinal relaxation time of blood spins (because blood is flowing to replenish fresh water spins) to save time, e.g., a TR of 15 ms was used for PC in mice at 11.7T scanner ^21^. Thus, partial volume effect is often controlled to be a small effect. By contrast, improper spatial resolution and slice positioning both lead to overestimation. Collectively, PC MRI tends to suffer from overestimation unless experimental parameters have been carefully optimized.

Slice positioning imperfection can be minimized by carefully placing the imaging slice by reference to the vascular trajectory ^32^. In order to improve the performance of slice positioning, time-of-flight (TOF) MRI can be utilized to track the trajectory of feeding arteries from different orientations, e.g., coronal and sagittal orientations ^21^. Taking a vessel with the diameter of 300 µm as an example, the maximal tolerable imperfection angle is 13.97°, showing agreement with an optimization study for human PC MRI where the tolerable imperfection angle is 15° ^33^.

CBF plays a central role in the microvascular functions and offers a sensitive marker for natural aging ^34^ and various diseases ^35-39^. For example, the CBF recovery pattern at the acute stage (within 3h post return of spontaneous circulation) can predict 24h neurological outcome in cardiac arrest ^40,41^. Apart from marking the pathological status, CBF can serve as an index for denoting pharmaceutical toxicity by tracking the extent that blood flow has been affected after drug delivery. Moreover, CBF is a fundamental parameter to calculate other higher-order physiological parameters, e.g., together with oxygen extraction fraction (OEF) to calculate cerebral metabolic rate of oxygen (CMRO2) ^42-44^, together with a vasoactive challenge to estimate cerebrovascular reactivity (CVR) ^24,25^, together with water extraction to calculate global BBB permeability ^27,28^. Therefore, CBF constitutes a popular and useful functional parameter to reveal the quantitative pathophysiology underlying diseases, where imaging and pathological assessments are often combined to provide a multiscale insight into the pathological mechanisms ^45-50^.

Findings in the current study should be interpreted with limitations. Different experimental parameters were simulated in a sequential order under the assumption that there was no interplay. It could be possible that these parameters affect each other interactively. Note that simulating several parameters simultaneously will lead to a hyperdimensional issue, making the results difficult to interpret. Projection to a lower dimension can simplify the issue to focus on major effects. In case the interplay between certain parameters is of interest, dedicated simulations can be designed and performed accordingly. Simulation as used in the current study facilitates the understanding of measurement accuracy for different techniques. A similar simulation study was performed to investigate the measurement accuracy of MR relaxation times ^51^. These simulations may serve as helpful guidance for *in-vivo* experiments.

## Conclusion

Accurate blood flow measurement can be achieved by phase contrast MRI when the following conditions are met: (i) spatial resolution is set to have at least six voxels along the vessel diameter; (ii) slice imperfection angle is less than 15° for a vessel diameter of 300 µm; (iii) normalized noise is less than 5.3%. Partial volume effect will underestimate the blood flow. These guidelines may facilitate the implementation of PC MRI in future pathophysiological studies.

## Acknowledgements

The author would like to thank Dr. Zhiliang Wei for proofreading and editing.

